# Convergent strigolactone perception via independent horizontal gene transfer of pre-adapted α/β-hydrolases

**DOI:** 10.1101/2025.03.14.642314

**Authors:** Qia Wang, Ye Ye, Lulu Wang, Yanlong Guan, Shuanghua Wang, Zhe Wang, Hang Sun, Steven M. Smith, Jinling Huang

## Abstract

Strigolactones (SLs) are not only phytohormones that influence multiple aspects of plant growth and development, but also signaling molecules for interactions between plants and certain fungi or bacteria. In plants, the SL receptor is an α/β-hydrolase (ABH) encoded by the *D14*/*KAI2* gene family, which is known to be derived from proteobacterial *RsbQ* through horizontal gene transfer (HGT). Here we show that the *CpD14*-*like* (*CDL*) gene family, which encodes another ABH exhibiting SL receptor functions in fungi, was acquired from actinobacteria via an independent HGT event. X-ray crystallographic experiments reveal that CDL and D14/KAI2/RsbQ (DKR) proteins have distinctive lid structures but maintain the same core ‘α/β fold’ and catalytic triad. Biochemical assays further show that both actinobacterial CDL and proteobacterial RsbQ can recognize and hydrolyze SLs, suggesting that they are pre-adapted for SL perception. This work demonstrates that independent HGT of pre-adapted genes can trigger convergent evolution of key innovations and provides insights into how SLs are sensed by organisms across kingdoms.

## Introduction

Strigolactones (SLs) are carotenoid-derived butenolides and are known to be important phytohormones. As endogenous hormones, SLs influence many aspects of plant growth and development, such as shoot branching, root architecture, and leaf senescence (1–3). SLs exuded by plants can also act as rhizosphere signals to stimulate seed germination of parasitic plants (4). Interestingly, an increasing number of studies indicate that SLs are also bioactive molecules sensed by certain fungi and bacteria, which in turn facilitate their interactions with plants (5). For instance, SLs can induce responses from arbuscular mycorrhizal (AM) fungi, such as spore germination and hyphal branching, thus promoting AM symbiosis (5, 6). Moreover, SLs have a widespread effect on the growth pattern of phytopathogenic fungi (7, 8). In bacteria, although the roles of SLs in root nodule symbiosis remain unclear, it has been reported that SLs could stimulate the swarming motility of rhizobacteria such as *Rhizobium leguminosarum* (9–11). These observations raise some intriguing questions: How can SLs act as signaling molecules perceived by various organisms across kingdoms? Is there evolutionary relatedness between the molecular mechanisms of SL perception in plants, fungi, and bacteria?

The molecular mechanisms of SL perception and signal transduction have been extensively studied in plants. An α/β-hydrolase (ABH) in seed plants named DWARF14 (D14) has been identified as the SL receptor (3, 12). D14 proteins have dual functions to sense and hydrolyze SLs, and their enzymatic activity against SLs requires an intact Ser/His/Asp catalytic triad (12–14). The *D14* genes arose from *KARRIKIN INSENSITIVE2* (*KAI2*) in seed plants through gene duplication and neo-functionalization (15). KAI2 proteins are receptors for exogenous karrikins (KARs), which are molecules present in smoke derived from burning plants and share a similar butenolide moiety with SLs (16, 17). The KAI2 signaling pathway also influences several developmental and physiological processes, such as seedling photomorphogenesis, leaf development, and abiotic stress tolerance (17, 18). Our previous study suggests that the plant *D14*/*KAI2* gene family was derived from proteobacterial *RsbQ* through horizontal gene transfer (HGT) (19). The acquisition of *RsbQ* not only was a critical step for the establishment of a butenolide sensing system (subsequently including SL signaling) in plants, but also represented a key innovation that facilitated plant adaptation to terrestrial environments.

Although SLs are also perceived by certain fungi and bacteria, the underlying mechanisms remain largely unclear. Recently, another ABH in the phytopathogenic fungus *Cryphonectria parasitica*, namely CpD14, was found to was found to be required for SL perception and influence fungal growth (20). CpD14 is similar to plant D14 in its ability to both bind and hydrolyze SLs, and such enzymatic activities also require an intact catalytic triad (12, 20). The *CpD14* knockout mutants of *C*. *parasitica* significantly reduced their sensitivity to synthetic SLs (stereospecific GR24); on the other hand, the SL receptor inhibitor tolfenamic acid alleviated the GR24 effect on the wild-type growth but had no impact on the mutants (20, 21). These observations are consistent with CpD14 serving as a receptor in an SL signaling pathway.

Here we provide evidence that the *CpD14-like* (*CDL*) gene family present in *C*. *parasitica* and closely related fungal groups was independently acquired from bacteria. Based on the structural and biochemical data, we reveal that HGT of bacterial ABHs can trigger convergent SL perception between plants and fungi through their conserved core protein structure and flexible substrate binding domain. These findings provide key insights into the origin and evolution of SL perception in different eukaryotic lineages.

## Results and discussion

### The fungal *CpD14*-*like* gene family has an actinobacterial origin

Fungal CpD14 and plant D14/KAI2 have only about 25% identity in their amino acid sequences, and they belong to different sub-clades of the ABH superfamily (22). Previous analyses identified CpD14 homologs in three fungal lineages, including ascomycetes, basidiomycetes, and Mucoromycota (20). To further assess the taxonomic distribution of CpD14 homologs, we performed extensive BLAST searches against available eukaryotic and prokaryotic genome sequence data (see Materials and Methods for details). We found that CpD14 homologs (E-value cutoff of 1e-6) are widely distributed across the Tree of Life (Data S1). Surprisingly, CpD14 homologs with significant sequence similarities (identity ≥ 40%) were overwhelmingly identified in the fungal group Leotiomyceta and the bacterial group actinobacteria (Fig. 1a). We named the hydrolase family of CpD14 and its significant homologs as CpD14-like (CDL). Aside from Leotiomyceta and actinobacteria, CDLs were identified in only a few species that scattered among different eukaryotic and prokaryotic lineages (Fig. 1a). Phylogenetic analyses showed that abnormally distributed CDLs from the same lineage (e.g., plants or basidiomycetes) did not form a monophyletic group, but were closely related to different Leotiomyceta or actinobacterial sequences (Fig. 1b), suggesting that they were derived from sequencing contamination or independent HGT events.

**Fig. 1.**
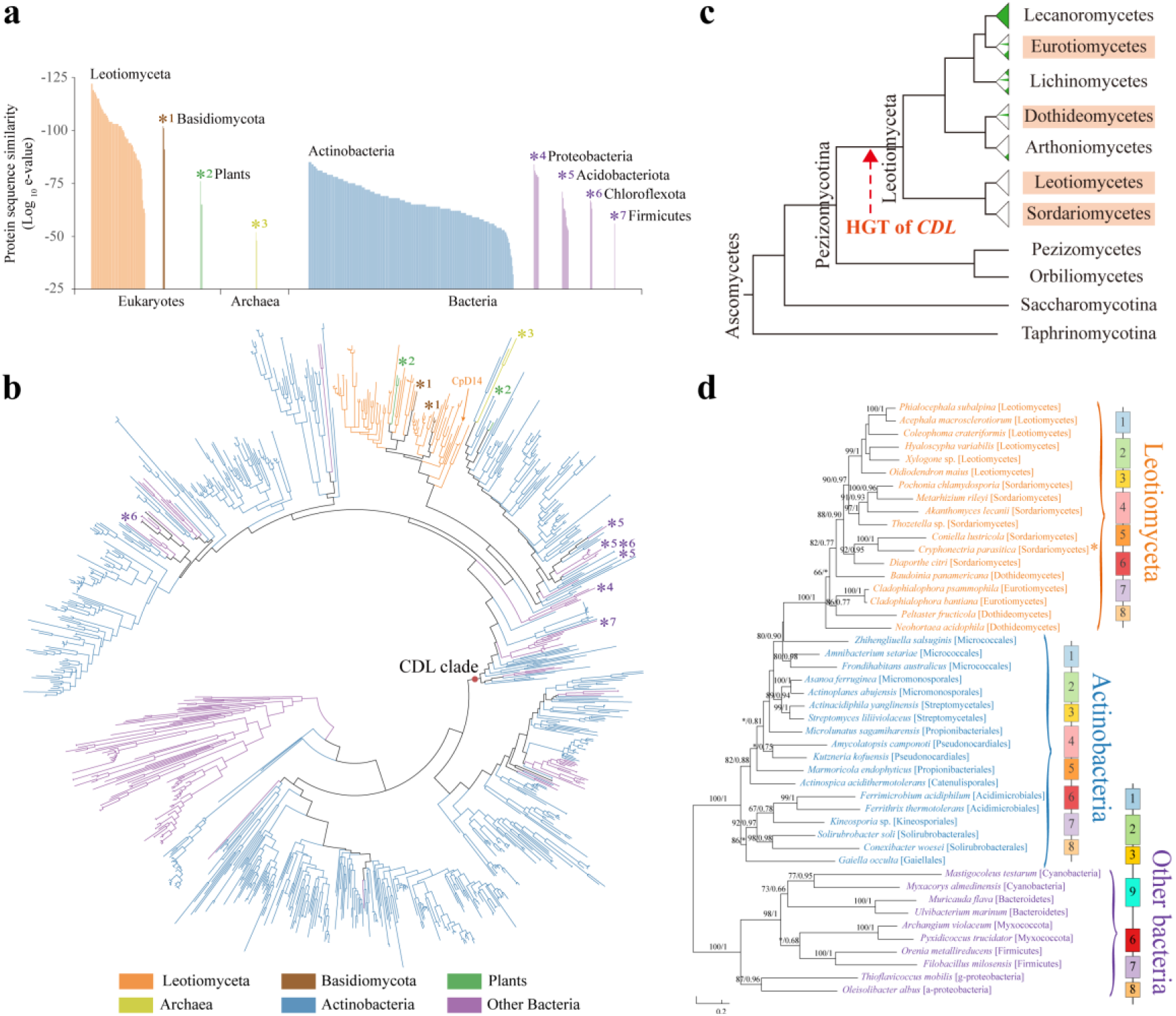
Fungal *CpD14-like* (*CDL*) gene family was derived from actinobacteria. **a**, Histogram of BLASTp E-values for CDLs in NCBI non-redundant (nr) protein sequences database (see additional details in Data S1). **b**, Phylogenetic analyses of CpD14 homologs performed with a maximum likelihood method using IQ-TREE. The abnormally distributed CDLs annotated in **a** are labeled with asterisks, and the node of CDL clade is labeled with a red dot. **c**, The evolutionary mode of *CDL*s in fungi. Green segments of the triangle indicate lichenized lineages, and fungal clades containing *CDLs* are highlighted in orange. **d**, Phylogenetic analyses of Leotiomyceta and actinobacterial CDLs. CpD14 was marked with an asterisk. CpD14 homologs from other bacteria were used as an outgroup. Numbers on branches show ultrafast bootstrap values (≥ 65%) from maximum likelihood method using IQ-TREE and posterior probabilities (≥ 0.65) from Bayesian method using MrBayes. Conserved motifs identified using MEME suite are indicated to the right (see details in Fig. S2).

The disjunct distribution of CDL sequences between Leotiomyceta and actinobacteria could theoretically be attributable to HGT between the two groups or vertical transfer followed by gene losses from all other groups (23, 24). We performed further analyses to distinguish these evolutionary scenarios. Using maximum likelihood and Bayesian methods, our phylogenetic analyses strongly supported a monophyletic group of Leotiomyceta CDLs, which in turn was nested within actinobacterial sequences (Fig. 1d). Sequence comparisons showed that CDLs of Leotiomyceta and actinobacteria share many highly conserved residues, including an identical catalytic triad (Fig. S1), as well as identical motif composition and organization (Fig. 1d; Fig. S2). The gene loss scenario not only requires numerous CDL loss events in all other eukaryotic and prokaryotic lineages, but also is incompatible with the high sequence conservation between the two distantly related lineages (i.e., Leotiomyceta and actinobacteria). We further estimated the divergence time of Leotiomyceta and actinobacterial CDL clade and found it to be less than 900 million years ago (mya) (Fig. S3). This divergence time is not only much younger than the split of eukaryotes (i.e., Leotiomyceta here) and prokaryotes (i.e., actinobacteria here) but also less than the age of the fungal kingdom (25), which argues strongly against the gene loss scenario. By contrast, HGT between Leotiomyceta and actinobacteria provides a straightforward and parsimonious explanation for the above relationship. Given the much earlier origin of actinobacteria than Leotiomyceta (25) and the presence of CpD14 homologs in other bacterial lineages (Fig. 1b, d), we reason that the *CDL* gene most likely evolved in bacteria and was then transferred from actinobacteria into Leotiomyceta.

### Structural insights into the convergent SL perception of different ABHs

Although varying in protein sequences, members of the ABH superfamily are characterized by a core ‘α/β fold’ domain that contains a nucleophile-his-acid catalytic triad, commonly Ser/His/Asp (26, 27) (Fig. 2a). The substrate selectivity of ABHs generally depends on an additional structural element, the lid domain (also known as cap domain), which regulates the accessibility of substrate binding crevices (26, 27). All D14/KAI2/RsbQ (DKR) structures reported in plants and bacteria so far have four additional α-helices (αT1-αT4), which form a V-shaped lid covering the ligand-binding pocket (28, 29) (Fig. 2a, b, d). Based on protein homology-modelling, previous studies speculated that the overall structure of CpD14 is similar to DKRs (20). We found that CDLs (including CpD14) and DKRs share identical residues in the catalytic triad and several critical folding sites, while CDLs typically have 24-26 extra residues in the lid domain (Fig. S4). These observations suggest the CDLs may be structurally similar to DKRs in the core ‘α/β fold’, but not in the lid part. However, no CDL structural data is currently available to confirm the above suggestion.

**Fig. 2.**
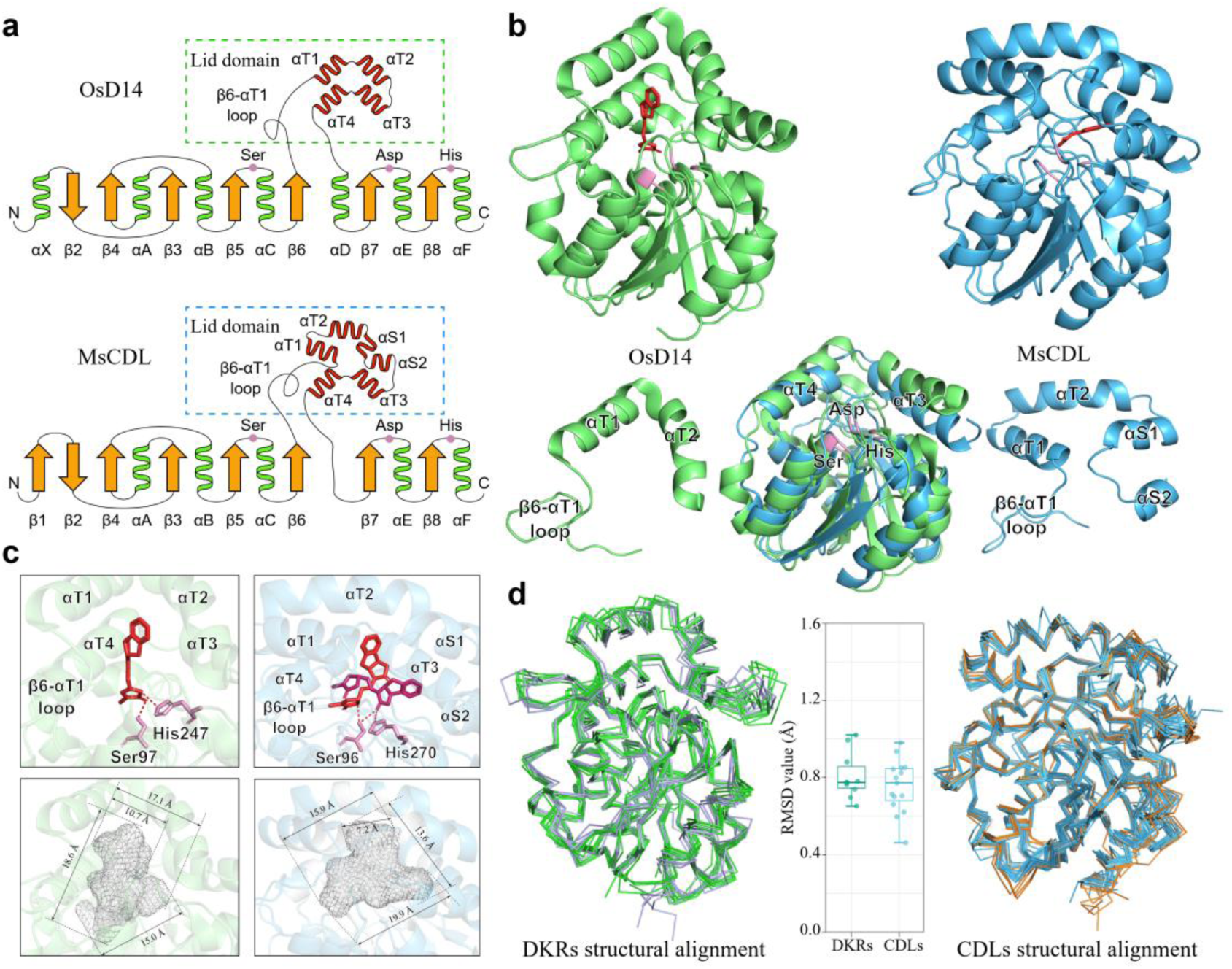
Structural characterization and comparison of D14/KAI2/RsbQ (DKR) and CpD14-like (CDL) proteins. **a**, The typical topology of DKR (represented by *Oryza sativa* D14) and CDL (represented by *Microlunatus sagamiharensis* CDL). **b**, The previously reported structure of OsD14 in complex with the synthetic SL GR24 (70) (PDB ID 5DJ5), and our determined structure of MsCDL in complex with 4-hydroxyphenylacetic acid (4HP). The 4HP was derived from the crystallization buffer and was observed in the ligand-binding pocket of MsCDL (linked with Ser and His residues of the catalytic triad). The GR24 and 4HP are in red, and the catalytic triads of OsD14 and MsCDL are in pink. The structural conservation and distinction between OsD14 and MsCDL are shown below. **c**, Binding configuration of GR24 in OsD14 (left, based on previously reported structural data (70)) and MsCDL (right, based on virtual docking data). Red dashed lines indicate the hydrogen or ionic bonds between ligands and residues of the catalytic triad. The size data of ligand-binding pockets are shown below. **d**, Structural alignments of different DKRs and CDLs. Structural alignments of plant D14/KAI2 (in green) and bacterial RsbQ (in purple) with OsD14 (in gray) are shown on left. Protein structures used in the analyses were downloaded from PDB (see Materials and Methods for PDB IDs). Structural alignments of the Alphafold2 (30) predicted structures of CDLs from different groups of Leotiomyceta (in orange, including CpD14) and actinobacteria (in blue) with MsCDL (in gray) are shown on right. The RMSD values of structural alignments are shown in the middle.

We therefore performed X-ray crystallographic experiments to determine the structure of a representative CDL from the actinobacterium *Microlunatus sagamiharensis* (termed here MsCDL). After recombinant protein expression and purification, MsCDL proteins were crystallized and X-ray diffraction data were collected to build an initial model, followed by multiple rounds of refinement (Table S1). Eventually, a high-resolution crystal structure (1.90 Å) of MsCDL was determined (Fig. 2b). As expected, MsCDL and *Oryza sativa* D14 (OsD14, as a representative of DKRs) share a conserved core ‘α/β fold’ structure, including spatial positions of the catalytic triad (Fig. 2b). Strikingly, unlike OsD14, the lid structure of MsCDL contains two additional short α-helices (αS1-αS2), and its αT1-αT2 α-helices turn about 90 degrees to form a triangular lid upper part (Fig. 2b). Nevertheless, the ligand-binding pocket of MsCDL resembles that of OsD14, and molecular docking simulations show that a canonical SL molecule can pass through the lid structure and bind in the ligand-binding pocket of MsCDL at different angles (Fig. 2c).

Using Alphafold2 (30), we predicted the structures of CDLs from different sub-groups of actinobacteria and Leotiomyceta (Fig. S5). Results of the predicted local-distance difference test (pLDDT) and predicted aligned error (PAE) showed that these predictions were accurate and reliable (Fig. S6). We then compared the structures of these predicted CDLs with the experimentally determined MsCDL and found that they are highly conserved (RMSD ≤ 0.98 Å; Fig. 2d; Fig. S5). We also performed structural predictions on RsbQ from different proteobacterial sub-groups and, together with the structure of *Bacillus subtilis* RsbQ determined (29), further confirmed the structural conservation between bacterial RsbQ and plant D14/KAI2 (RMSD ≤ 1.02 Å; Fig. 2d; Fig. S5). Taken together, the above data indicate that the Leotiomyceta CDL and plant D14/KAI2 belong to two distinct structural types of ABHs (i.e., CDL and DKR types) inherited from different bacterial lineages, representing convergent evolution in molecular functions.

In addition to CDLs and DKRs, *Arabidopsis* carboxylesterases (CXEs, including AtCXE15 and AtCXE20, also belonging to ABHs) have also been found to bind and/or hydrolyze SLs, even though they are more likely to act as hydrolases in SL catabolism, rather than as receptors in SL perception (31, 32). The structures of AtCXE15 and AtCXE20 were elucidated by a recent study (33) and their lid structures are significantly different from those of CDLs or DKRs (Fig. S7), again suggesting that the lids can be relatively flexible for SL binding and hydrolysis. Notably, we observed that multiple actinobacterial CDLs are annotated as pimeloyl-ACP methyl ester CXEs (Data S1). In bacteria, various other ABHs (e.g., BioH, BioV, BioG, BioK, and BioJ) are also recruited to function as pimeloyl-ACP methyl ester CXEs (34). Similar to CDLs and DKRs, these CXEs belong to different sub-clades of the ABH superfamily but catalyze the same carboxylesterase reaction in biotin (vitamin H) biosynthesis (35, 36). Structural comparisons of these CXEs showed that some of them (e.g., BioG and BioH) are similar to DKRs, while others (e.g., BioJ) are similar to AtCXEs (Fig. S7). These additional data show that it may be common for ABHs with varying lid structures to catalyze the same reaction on similar substrates.

### Bacterial ABHs were pre-adapted for SL perception

To better understand the origin of SL perception in ABHs, we performed additional experiments to investigate the biochemical properties of actinobacterial CDL and proteobacterial RsbQ. The recombinant proteins of MsCDL and proteobacterial *Stenotrophomonas rhizophila* RsbQ (SrRsbQ) were expressed and purified for experiments, and fungal CpD14 and plant *Arabidopsis thaliana* D14 (AtD14) were also prepared for comparisons. Subsequently, we investigated the binding and hydrolytic activity of these ABH proteins toward different types of SLs, including Yoshimulactone green (YLG), the canonical SL (with an intact ABC tricyclic lactone) analog GR24 and its desmethyl equivalent dGR24, as well as the non-canonical SL (lacking the A, B, or C ring) analog 4-Br debranone (4BD) and its desmethyl equivalent d4BD (Fig. 3a).

**Fig. 3.**
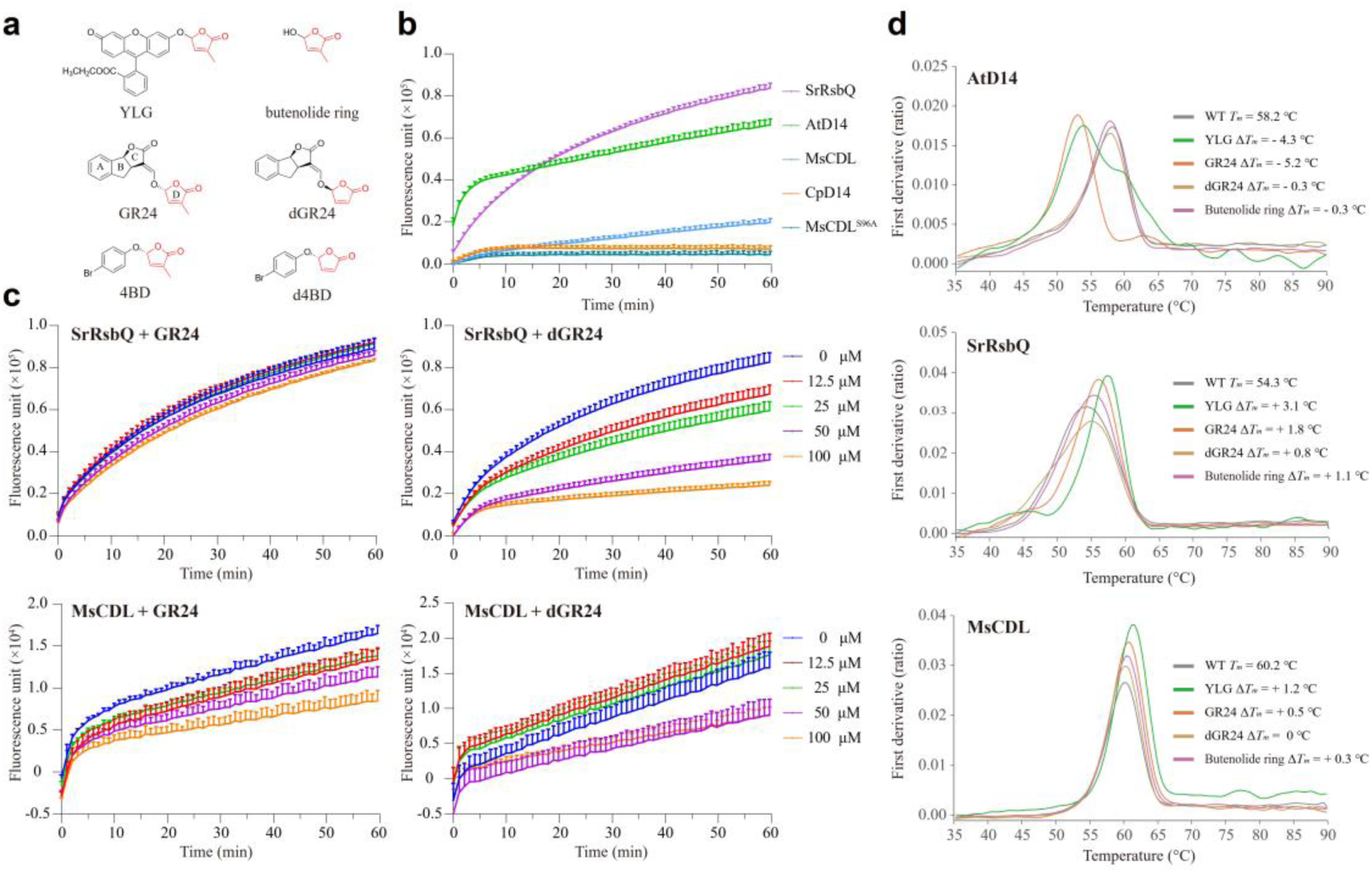
Binding and hydrolytic activity of DKRs and CDLs toward SL analogs. **a**, The structures of SL analogs and butenolide ring used in this study, with the butenolide moiety highlighted in red. GR24, dGR24, 4BD, and d4BD are all equimolar racemic mixtures of two enantiomers. **b**, Real-time monitoring YLG hydrolysis assays. Data are means ± s.d.; *n* = 3 technical replicates. **c**, YLG hydrolysis assays in the presence of GR24 and dGR24, respectively. The concentration gradients of GR24 and dGR24 are shown on the right. Data are means ± s.d.; *n* = 3 technical replicates. **d**, Thermal shift assays. The melting temperature curves represent the means of 3 technical replicates. The melting temperature (*T*_m_) of protein and its variations (Δ*T*_m_) after 30 min incubation with substrates are shown on the right of each group.

We first examined the hydrolytic activity of these proteins toward YLG, a fluorescent SL probe whose fluorescence emission could be activated once hydrolysed (37). In real-time monitoring experiments, SrRsbQ and MsCDL exhibited higher YLG hydrolytic activities than AtD14 and CpD14, respectively (Fig. 3b). Substitution of the catalytic residue S96A in MsCDL (MsCDL^S96A^) significantly reduced but not completely abolished its hydrolytic activity towards YLG, as was also reported for RsbQ from the firmicute *Bacillus subtilis* (38). Moreover, the non-fluorescent SL analogs dGR24 and GR24 can inhibit the YLG hydrolytic activities of SrRsbQ and MsCDL in a dose-dependent manner, among which the inhibition of SrRsbQ by dGR24 is the most significant (Fig. 3c). These data indicate that both bacterial RsbQ and CDL are not only efficient SL hydrolases (their hydrolytic activities also rely on the catalytic triad), but also can bind different types of SLs.

We further used the LC-MS method to examine the binding activity of the above ABHs toward SLs. After one hour of incubation with YLG or GR24, a mass increase peak of +96 Da corresponding to the butenolide moiety was detected in the intact denatured mass spectrometry of AtD14, SrRsbQ, and CpD14 (Fig. 4a). In comparison, no such a peak was detected when these proteins were directly incubated with the butenolide ring, suggesting that the tested ABHs could recognize the intact SL molecules and bind their butenolide moiety after hydrolysis. Moreover, the results of digested protein mass spectrometry confirmed that the butenolide moiety was covalently linked to the catalytic histidine residue of SrRsbQ (Fig. 4b), as observed in AtD14 (12). On the other hand, when AtD14, SrRsbQ, and MsCDL were incubated with dGR24, each yielded a peak of +284 Da corresponding to an intact dGR24 molecule (Fig. 4a). Similar binding patterns were also observed when these proteins were incubated with 4BD and d4BD (Fig. S8a). These observations further suggest that bacterial RsbQ and CDL can bind and hydrolyze SLs, but do not necessarily bind to the butenolide moiety after hydrolysis.

**Fig. 4.**
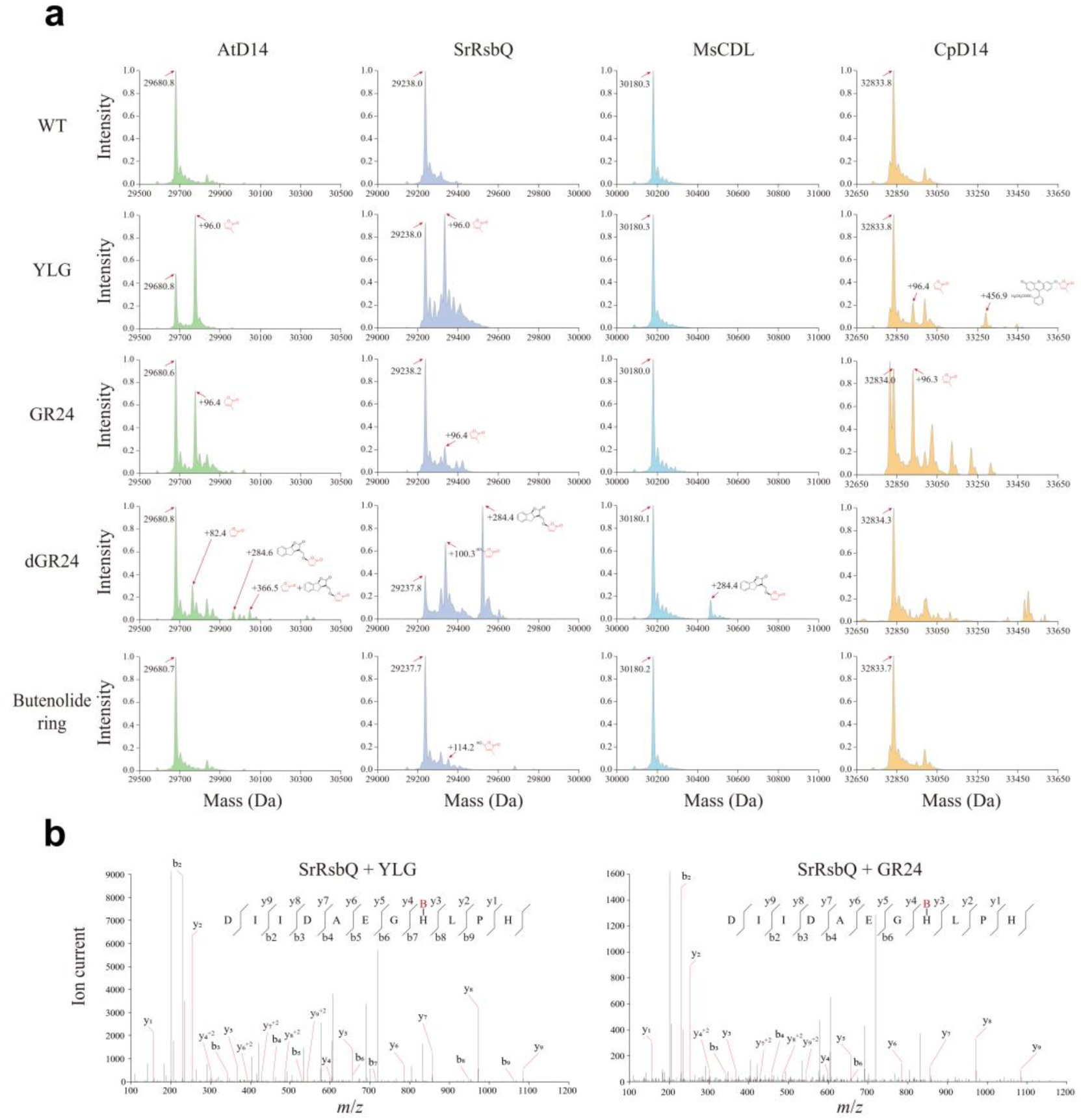
Mass spectrometry of DKRs and CDLs toward SL analogs. **a**, Deconvoluted intact mass spectra of DKRs and CDLs after 1 h incubation with SL analogs and butenolide ring, using no-substrate samples as controls (WT). The peaks of native proteins and their putative bound states are labeled with molecular weight values. **b**, Digested mass spectrometry of SrRsbQ treated with chymotrypsin digestion following YLG (left) and GR24 (right) incubation.

Ligand binding usually influences protein conformation, thereby changing its thermal stability and causing a shift in melting temperature (*T*_m_) (39). We therefore conducted thermal shift assays (TSA) to examine the conformational change of the above ABHs toward different SLs. For AtD14, both YLG and GR24 could result in a significant *T*_m_ variation (Δ*T*_m_ ≥ 4.3 ℃; Fig. 3d), which reflects a drastic conformational change in the lid domain that forms an interaction interface with signal transduction components (12). Due to the intrinsic nature of CpD14, its melting curve is unstable, precluding determination of the *T*_m_ value (Fig. S8b), as previously reported (20). Nearly all types of SLs could induce a *T*_m_ shift in SrRsbQ and MsCDL, however, more significant *T*_m_ variations (Δ*T*_m_ ≥ 1.8 ℃) were observed in those protein groups that bind to the butenolide moiety of SLs after hydrolysis (Fig. 3d; Fig. 4a).

Taken together, our data show that bacterial RsbQ and CDL have various degrees of affinity and hydrolytic activity for different types of SLs, suggesting that they are appreciably pre-adapted (or have the potential) SL perception. It has been known that the switch of substrate selectivity (or specificity) for many ABHs might result from simple substitutions in amino acid sequences. For instance, SL receptors evolved from *KAI2* genes more than once in plants (15, 40), and the conversion from KAI2 to an SL receptor required only three amino acid changes in the lid domain (41). In parasitic plant *Striga hermonthica*, members of the KAI2 family differ in relatively few amino acids but exhibit different substrate specificities, while single amino acid substitutions can significantly change its substrate specificity (42). Presumably, similar substitutions could ultimately transform the independently acquired and pre-adapted ABHs into specific SL receptors in plants and fungi.

### A model for the evolution of SL perception in organisms

Within Leotiomyceta, *CDL* genes were only found in four classes, including Eurotiomycetes, Dothideomycetes, Leotiomycetes, and Sordariomycetes (Fig. 1c). Recent phylogenomic studies showed that Eurotiomycetes, Dothideomycetes, and several predominantly lichenized classes form a clade, which in turn is sister to another clade comprising Leotiomycetes and Sordariomycetes (43, 44). Therefore, we speculate that the *CDL* gene was initially acquired by the last common ancestor of Leotiomyceta about 450 mya, and subsequently subjected to secondary losses in other descendent groups (Fig. 1c). Intriguingly, we found that the acquired *CDL* gene is now retained in pathogenic, endophytic, or saprophytic fungi, but lost in almost all lichenized lineages in Leotiomyceta (with the only exception of *Viridothelium virens*, see Data S2).

Previously, a gene encoding methylammonium permease (MEP) was also reportedly transferred from prokaryotes to the last common ancestor of Leotiomyceta; however, the *MEP* gene has been preferentially retained in lichenized fungi (45). It was suggested that MEP may contribute to nitrogen transport between lichenized fungi and their algal partners in lichen symbioses (45). Because the diversification of Leotiomyceta was accompanied by various forms of interaction (pathogenic, parasitic, or endophytic) with land plants (46), we reason that the opposite retention/loss pattern of *CDL* to *MEP* may also be related to its specific role in plant-fungal interactions. Recent studies suggest that SL biosynthesis evolved in land plants about 500 mya (47), and SLs initially served as symbiotic rhizosphere signals between land plants and fungi rather than endogenous hormones (48). Given that the HGT of *CDL* gene in Leotiomyceta followed closely after the origin of SL biosynthesis in land plants (Fig 5), it is likely that, immediately after its acquisition, *CDL* was coopted by the Leotiomyceta to perceive land plant SLs, thereby facilitating their interactions. This assumption might explain the observation that *CDL* is now preferentially retained in those Leotiomyceta fungi interacting with land plants.

**Fig. 5.**
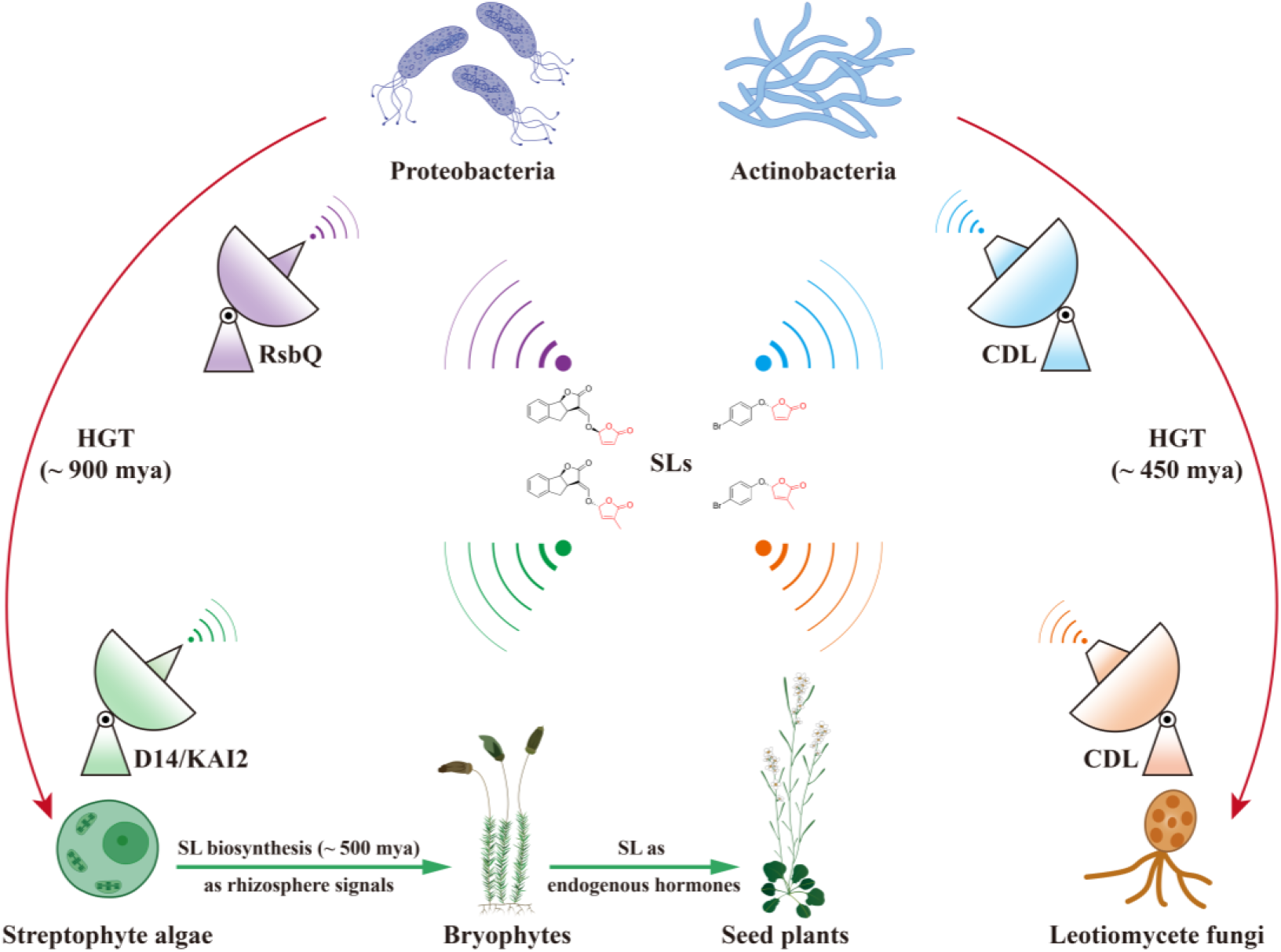
A proposed model for the evolutionary trajectory of SL perception. Before SLs were produced by plants, bacteria had already evolved several types of ABHs (including RsbQ and CDL) to cleavage butenolides or similar esters/lactones. The basic structural framework of these ABHs determines that they have the potential to be SL receptors (we characterized these pre-adapted receptors as radars receiving SL signals). About 900 million years ago (mya), the proteobacterial RsbQ was acquired by the ancestor of streptophytes via a HGT event, and turned into plant D14/KAI2 to catalyze similar esters/lactones cleavage reaction (19). During streptophyte evolution, SL biosynthesis first evolved in land plants at about 500 mya through the recruitment of *D27*, *CCD7*, and *CCD8* genes which were initially originating from cyanobacteria via endosymbiotic gene transfer (EGT) (19). Subsequently, D14/KAI2 were gradually recruited for SL signaling systems as receptors (47). In fungi, the actinobacterial *CDL* gene was acquired by the ancestor of Leotiomyceta fungi via an independent HGT event about 450 mya. Leotiomycete fungi may coopt CDL to perceive land plant SLs and facilitate their interactions right after the acquisition of CDL, which is consistent with the initial role of SLs as symbiotic rhizosphere signals in land plants (48). In summary, the formation of SL signaling system in organisms was accompanied with the acquirement and integration of genetic resources from different sources.

Combined with the current knowledge, we propose the following model to describe the evolutionary trajectory of SL perception (Fig. 5). Many esters and lactones (cyclic esters), including butenolides, are produced by plants, fungi, and bacteria through the carotenoid or other isoprenoid biosynthetic pathways (49, 50). It is likely that bacteria first evolved various ABHs to hydrolyze these esters and lactones. Some of these bacterial ABH genes, such as *RsbQ* and *CDL*, were spread to plants and fungi via independent HGT events. Due to their substrate promiscuity, these acquired ABHs may immediately catalyze the cleavage reaction upon some endogenous or exogenous butenolides (or similar esters/lactones), thus providing an adaptive advantage to the recipient lineages. Over time, these structurally and functionally pre-adapted ABHs can evolve into specific types of SL receptors in plants, fungi, and even bacteria through simple amino acid substitutions. Such a convergent SL perception builds a signal system between plants and microbes, thus promoting their interactions. The SL receptors in AM fungi have not yet been identified, and no significant homologs of CDLs or DKRs were found in their genomes. We thus reason that AM fungi may recruit another type of ABH as SL receptor. Given the widespread distribution of ABHs and the less strict structural requirements for SL receptors, independent evolution of SL perception in organisms may occur more frequently than we have realized.

## Conclusions

In this study, we provide evidence that the Leotiomyceta *CDL* gene family for SL perception was derived from actinobacteria via HGT. Together with our previous finding that the *D14*/*KAI2* gene family in plants was acquired from proteobacteria (*19*), it is clear that in both plants and fungi, the ABHs currently known to be associated with SL perception are of bacterial ancestry. We suggest that some bacterial ABHs are pre-adapted for SL perception by possessing a basic structural framework, but they are structurally flexible to recognize various substrates in different metabolic pathways. The acquisition of these ABH genes might have initially provided plants and fungi with the same or similar metabolic functions, and subsequent changes (e.g., simple amino acid substitutions) in these hydrolases led to the convergent SL perception. This work not only demonstrates that independent HGT events of pre-adapted genes can trigger convergent evolution of major innovations, but also provides insights into the origins of SL perception in different organismal groups across kingdoms.

## Materials and Methods

### Identification of CpD14 homologs

The protein sequence of *Cryphonectria parasitica* D14 (CpD14, NCBI accession number XP_040779045.1) was used as a query for BLAST searches against NCBI non-redundant (nr) protein sequences database (https://www.ncbi.nlm.nih.gov/). An E-value cutoff of 1e-6 was applied. The distribution of CpD14 homologs in eukaryotes was further verified by BLAST searches against other sequence databases, including the 1,000 Plants Project (OneKP, https://db.cngb.org/onekp) (51), the JGI Fungal Program (https://mycocosm.jgi.doe.gov/mycocosm/home) (52), and the Marine Microbial Eukaryote Transcriptome Sequencing Project (MMETSP, https://www.imicrobe.us/#/projects/104) (53). The search results are listed in Data S1.

### Phylogenetic analyses

Multiple alignments of protein sequences were performed using MAFFT v7.520 (54) and trimmed using trimAl v1.4 (55) with the parameter “-automated1”. Maximum likelihood phylogenetic analyses were performed using IQ-TREE v2.2.0 (56) with 1,000 replicates for both ultrafast bootstrap and SH approximate likelihood ratio test. Bayesian phylogenetic analyses were performed using MrBayes v3.2.7 (57) with two independent runs each of two million generations, sampled every 1,000 generations, and the first 25% samples were discarded as burn-in. The best-fit substitution models were determined by ModelFinder (58). Phylogenetic trees were visualized using FigTree v1.4.4 (http://tree.bio.ed.ac.uk/software/figtree/).

### Protein sequence conservation analyses

Protein sequence alignments were performed using MAFFT v7.520 (54). Visualization of conserved amino acid residues and annotation of secondary structures were performed using ESPript 3.0 (59). The global similarity score that defines the threshold of high similarity was set to 0.7. Conserved motifs were identified using Multiple Em for Motif Elicitation (MEME) suite 5.4.1 (https://meme-suite.org/meme/tools/meme) (60). Motif E-value threshold was set to 1e-8. Visualization of motifs was performed using TBtools v2.031 (61).

### Divergence time estimation

The divergence time estimation was performed using BEAST v2.7.4 (62). Gamma category count was set to 4, and the best-fit substitution models were determined by ModelFinder (58). The Calibrated Yule Model was adopted for the tree prior. According to previous studies (46), the node time of Leotiomyceta was set as 450 mya (mean time, with a sigma value of 20 mya) to calibrate nodes on the tree. Markov Chain Monte Carlo (MCMC) analyses were performed with ten million chains under the uncorrelated relaxed clock model to reach an effective sample size (ESS) of at least 200 in Tracer v1.7.2 (63). The maximum clade credibility (MCC) consensus tree was generated using TreeAnnotator v2.7.7 (62) after discarding the first 10% samples as burn-in, and visualized using FigTree v1.4.4 (http://tree.bio.ed.ac.uk/software/figtree/).

### Protein expression and purification

The N-terminal 6His-Strep Ⅱ-TEV fusion proteins of MsCDL, MsCDL^S96A^, and SrRsbQ, and the N-terminal 4T-3-GST-TEV fusion protein of CpD14 were each expressed in *Escherichia coli* strain BL21 (DE3). The N-terminal SUMO-Thrombin-6His-Strep Ⅱ-TEV fusion protein of AtD14 was expressed in *E. coli* strain Rosetta2 (DE3). *E. coli* cells were grown in LB media and induced with 0.1 mM isopropyl-β-D-thiogalactoside (IPTG) at 16 ℃ for 16 h. Subsequently, the cells were collected and resuspended with lysis buffer [50 mM Tris-HCl (pH 7.5 or 8.0), 500 mM NaCl, 10% (v/v) glycerol], and lysed 3-5 times with a high-pressure homogenizer at 750 bar. Lysates were centrifuged at 30,700 ×g for 30-60 min at 4 ℃. The supernatant was collected for affinity chromatography (His FF column for MsCDL, MsCDL^S96A^, SrRsbQ, and AtD14; Ni-NTA column for MsCDL and AtD14; Strep-Biortus column for MsCDL; GST column for CpD14; Strep-Tactin XT column for AtD14). Tags of all proteins were removed by incubation with TEV protease for 12 h in lysis buffer. The samples were further purified by size-exclusion chromatography (SEC; Superdex 200 increase 10/300 GL column for MsCDL; HiLoad 16/600 Superdex 200 pg column for MsCDL^S96A^, SrRsbQ, CpD14, and AtD14) and ion-exchange chromatography (Source 15Q column for CpD14). The eluted proteins were identified by SDS-PAGE, analytical SEC, and LC-MS. Proteins in HEPES buffer [20 mM HEPES (pH 7.5), 150 mM NaCl] were stored at -80 ℃ before being used for analyses.

### Crystallization, X-ray diffraction data collection, and structure determination

MsCDL protein sample (0.2 μL) was mixed with 0.18 μL crystallization solution [0.1 M Bis-Tris (pH 6.1), 1.8 M (NH_4_)_2_SO_4_ and 0.02 μL Silver Bullets (0.25% (w/v) 1,3,5-pentanetricarboxylic acid, 0.25% (w/v) 4-hydroxyphenylacetic acid, 0.25% (w/v) benzoic acid, 0.25% (w/v) poly (3-hydroxybutyric acid), and 0.02 M HEPES sodium (pH 6.8)] using the hanging drop vapor diffusion method. Crystals appeared in about 3 days at 20 ℃. Crystals with appropriate size (typically 10.6 μm × 281.6 μm) were collected and fast cooled in liquid nitrogen for subsequent data collection.

A crystal producing high-quality diffraction pattern was chosen for full dataset collection at the Canadian Light Source (CLS) of the University of Saskatchewan, Canada. The diffraction data were indexed and integrated with X-ray Detector Software (XDS) (64) and scaled by Aimless (65). The space group is P41212 and the resolution reached 1.90 Å. Crystal structure was solved by molecular replacement with PHASER (66). After multiple rounds of refinement with Coot (67) and Refmac5 (68), the final coordinate had *R*_work_ value of 14% and *R*_free_ value of 18%. Crystallographic parameters and the statistical data collection and refinement are shown in Table S1. The structural information of MsCDL has been uploaded to the Protein Data Bank (PDB) with an entry ID of 8YIJ.

### Protein structural prediction, comparison, and docking simulation

Tertiary structures of CDLs from representative groups of Leotiomyceta and actinobacteria, which have not been experimentally determined, were predicted by AlphaFold2 (30). The accuracy of predictions was assessed based on the predicted local-distance difference test (pLDDT) and predicted aligned error (PAE) scores (Fig. S6). The previously reported structures of D14/KAI2/RsbQ (DKRs) and carboxylesterases (CXEs) were downloaded from PDB, including entry IDs of 5DJ5 (*Oryza sativa* D14), 4IH4 (*Arabidopsis thaliana* D14, AtD14), 7TVW (AtD14-like 2), 4JYP (AtKAI2), 7UOC (*Orobanche minor* KAI2d4), 5Z7X (*Striga hermonthica* HTL4), 6AZB (*Physcomitrium patens* KAI2-like E), 1WOM (*Bacillus subtilis* RsbQ), 8VCD (AtCXE15), 8VCE (AtCXE20), 5GNG (*Haemophilus influenzae* BioG), 4ETW (*Shigella flexneri* BioH), and 6K1T (*Francisella philomiragia* BioJ). Protein structural alignment and RMSD calculation were performed with PyMOL (http://www.pymol.org/). Docking simulations of MsCDL protein with GR24 were performed using AutoDock v4.2.6 (69). The grid size of X, Y, and Z were each set as 60, and grid space as 0.375. Visualization of docking simulations was generated with PyMOL.

### Chemical compounds

The Yoshimulactone Green (YLG) and 5-Hydroxy-3-methyl-2(5H)-furanone (butenolide ring) were purchased from Tokyo Chemical Industry (TCI, Shanghai), and GR24 (rac-GR24, an equimolar racemic mixture of two enantiomers, GR24^5DS^ and GR24^ent-5DS^) was purchased from StrigoLab. The 4BD was provided by Ruifeng Yao (Hunan University, China), the same as described previously (12). The dGR24 and d4BD (both are equimolar racemic mixtures of two enantiomers) were synthesized and then verified by high resolution mass spectrometer (HR-MS) and nuclear magnetic resonance (NMR). For details of dGR24 and d4BD syntheses, see Fig. S9.

### YLG hydrolysis assays

The real-time monitoring YLG assay was conducted with 10 μM YLG and 0.25 μM proteins (final concentration) in 100 μL reaction buffer [100 mM HEPES (pH 7.4), 150 mM NaCl] in a 96-well black plate (OptiPlate). The fluorescence intensity was measured using a SpectraMax iD3 (Molecular Devices, San Jose) microplate reader. The measurements were performed at 28 ℃ with an excitation wavelength of 480 nm and an emission wavelength of 520 nm, Photomultiplier Tube (PMT) Gain of 500 volts, and read height of 2 mm. For non-fluorescent SL competition assays, YLG was premixed with 0-100 μM GR24 or dGR24 before proteins were added. The fluorescence intensity was calculated by subtracting the background hydrolysis in no-protein control groups.

### Intact denatured protein mass spectrometry

AtD14, SrRsbQ, CpD14, and MsCDL proteins were each added at a final concentration of 10 μM and incubated with 200 μM substrate in the reaction buffer [100 mM HEPES (pH 7.4), 150 mM NaCl] at 25 ℃ for 1 h, and centrifuged at 13,800 ×g for 5 min before detecting. Each group was repeated two times. The UPLC-Q-TOF system was constructed by a Shimadzu LC system controller coupling to a SCIEX 6600^+^ TripleTOF mass spectrometer. Chromatography was performed using a BioZen Intact C4 column (150 × 4.6 mm) with 3.6 μm particle size. The following LC conditions were used for the analyses: injection volume 5 μL; flow rate 0.6 mL min^-1^; column temperature 60 ℃; mobile phase included water + 0.1% (v/v) formic acid (Pump A) and acetonitrile + 0.1% (v/v) formic acid (Pump B); gradient elution program, 0-2 min 15% (v/v) acetonitrile/H_2_O, 2-7 min 15%-60% acetonitrile/H_2_O, 7-7.5 min 60%-90% acetonitrile/H_2_O, 7.5-10 min 90% acetonitrile/H_2_O, 10-10.5 min 90%-15% acetonitrile/H_2_O, 10.5-15 min 15% acetonitrile/H_2_O. The mass spectrum conditions were as follows: spray voltage 5500 V for the positive-ion mode; *m*/*z* range from 100 to 6000; gas temperature 500 ℃; curtain gas 35 psi; nebulizer gas 55 psi; auxiliary gas 55 psi; collision energy 10 V; declustering potential 140 V.

### Digested protein mass spectrometry

After 1 h incubation with 200 μM YLG or GR24, SrRsbQ proteins (20 μg) were concentrated and diluted with guanidine hydrochloride (3.5 M, final concentration) at 55 °C for 1h, then alkylated with iodoacetamide (12.5 mM, final concentration) at room temperature for 30 min in the dark. Before digestion, the guanidine hydrochloride concentration in solution was diluted to less than 1 M with 50 mM Tris-HCl, and then digestion was performed with chymotrypsin (Promega) at 25 °C for 15 h. The reaction was stopped by adding 2 μL formic acid. The solution was desalted through C18 Sep-Pak column (Waters), then dried and the samples were redissolved into 0.1% (v/v) formic acid.

Samples were analyzed by liquid chromatography-tandem mass spectrometry (LC-MS/MS) using a TripleTOF 6600 mass spectrometer (SCIEX, Framingham) with an Optiflow ion source, coupled with an Eksigent NanoLC 400 system (SCIEX, Framingham). Mobile phase A was 2% (v/v) acetonitrile (ACN, Merck), 0.1% (v/v) formic acid, and mobile phase B was 98% (v/v) ACN, 0.1% (v/v) formic acid. Peptides of SrRsbQ were enriched and washed in a trap column (5 μm, ChromXP C18CL, 120 Å, 10 × 0.3 mm) at a flow rate of 10 μL min^-1^ for 3 min by mobile phase A, and then separated in a column (3 μm, ChromXP C18CL, 120 Å, 150 × 0.3 mm) with 5 μL min^-1^ by increasing mobile phase B. Data were acquired in positive ion mode using a data-dependent strategy. The method consisted of TOF MS scan for precursor ions with *m*/*z* ranging from 350 to 2000, followed by MS/MS scans for fragment ions with *m*/*z* ranging from 350 to 2000, allowing for a maximum of 30 candidate ions being monitored per cycle (50 ms accumulation time, 50 ppm mass tolerance, rolling collision energy, charge state +2 to +5, exclusion of ions after two fragmentations for 30 s). LC-MS/MS data were analyzed against the SrRsbQ database by ProtenPilot software v5.0.2 (SCIEX), considering chymotrypsin digestion and several possible modifications of serine, aspartic acid, and histidine with +96.021120 (butenolide moiety).

### Thermal shift assays

AtD14, SrRsbQ, CpD14, and MsCDL proteins with a final concentration of 10 μM were each incubated with 200 μM substrate in 10 μL reaction buffer [100 mM HEPES (pH 7.4), 150 mM NaCl] at room temperature for 30 min. The thermal melting curve was monitored using Tycho NT.6 (NanoTemper, Munich) with a ramp rate of 30 ℃ min^-1^ from 35 ℃ to 95 ℃.

## Acknowledgments

We thank Jian Liu [Kunming Institute of Botany, Chinese Academy of Sciences (KIB, CAS), China] and Yongsheng Chen (Liaoning Normal University, China) for technical suggestions on divergence time analyses, Lian Yang (KIB, CAS, China), Hongmei Li (KIB, CAS, China), and Xiaolei Lv (Shanghai AB Sciex Analytical Instrument Trading Co., China) for technical assistance in biochemical experiments, Ruifeng Yao (Hunan University, China) for providing the chemical compound 4BD. This work is supported in part by grants from the National Natural Science Foundation of China (NSFC) (32170242, 32470244), the Youth Innovation Promotion Association CAS (2022398), the Yunnan Revitalization Talent Support Program "Young Talent" Project (XDYC-QNRC-2022-0046), the Yunnan Fundamental Research Projects (202201AT070163), the Second Tibetan Plateau Scientific Expedition and Research (STEP) Program (2019QZKK0502), the National Key R&D Program of China (2024YFF1306700), and the Australian Research Council Centre of Excellence for Plant Success in Nature and Agriculture (CE200100015).

## Author contributions

Q.W., S.M.S., and J.H. conceived and designed the study. Q.W., Y.Y., L.W., Y.G., S.W., Z.W., H.S., S.M.S., and J.H. performed the experiments and data analyses. Q.W., S.M.S., and J.H. wrote the manuscript, and all authors contributed to the manuscript editing.

## Extended Data

**Fig. S1.**
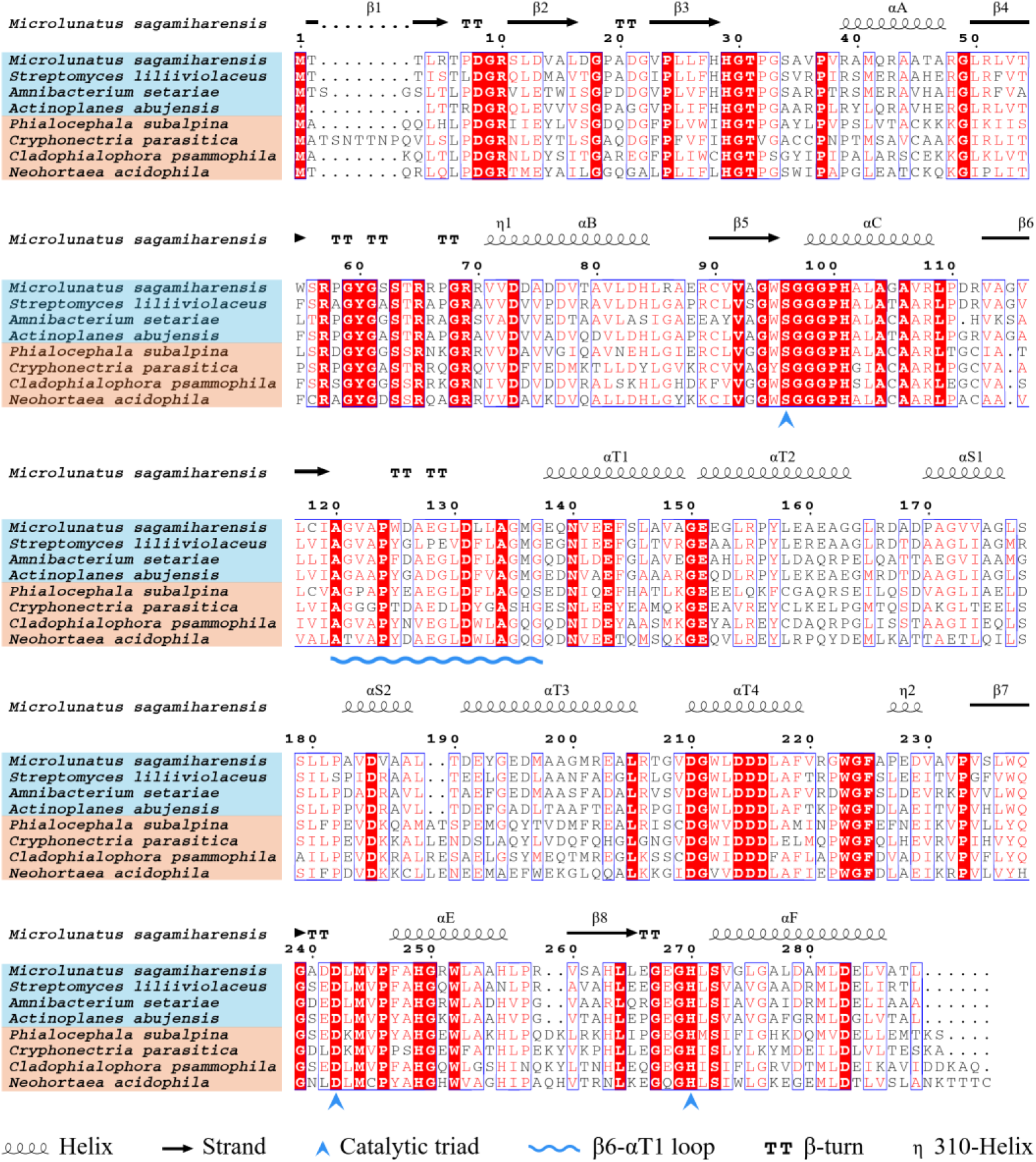
Sequence conservation between actinobacterial and Leotiomyceta CDLs. In the first column, actinobacterial CDLs are highlighted with a light blue, and Leotiomyceta CDLs are highlighted with a light brown frame. Secondary structural annotations based on our determined tertiary structure of MsCDL are displayed on top of sequences. Identical and highly conserved residues are highlighted with red and white rectangular boxes respectively, both framed in blue.

**Fig. S2.**
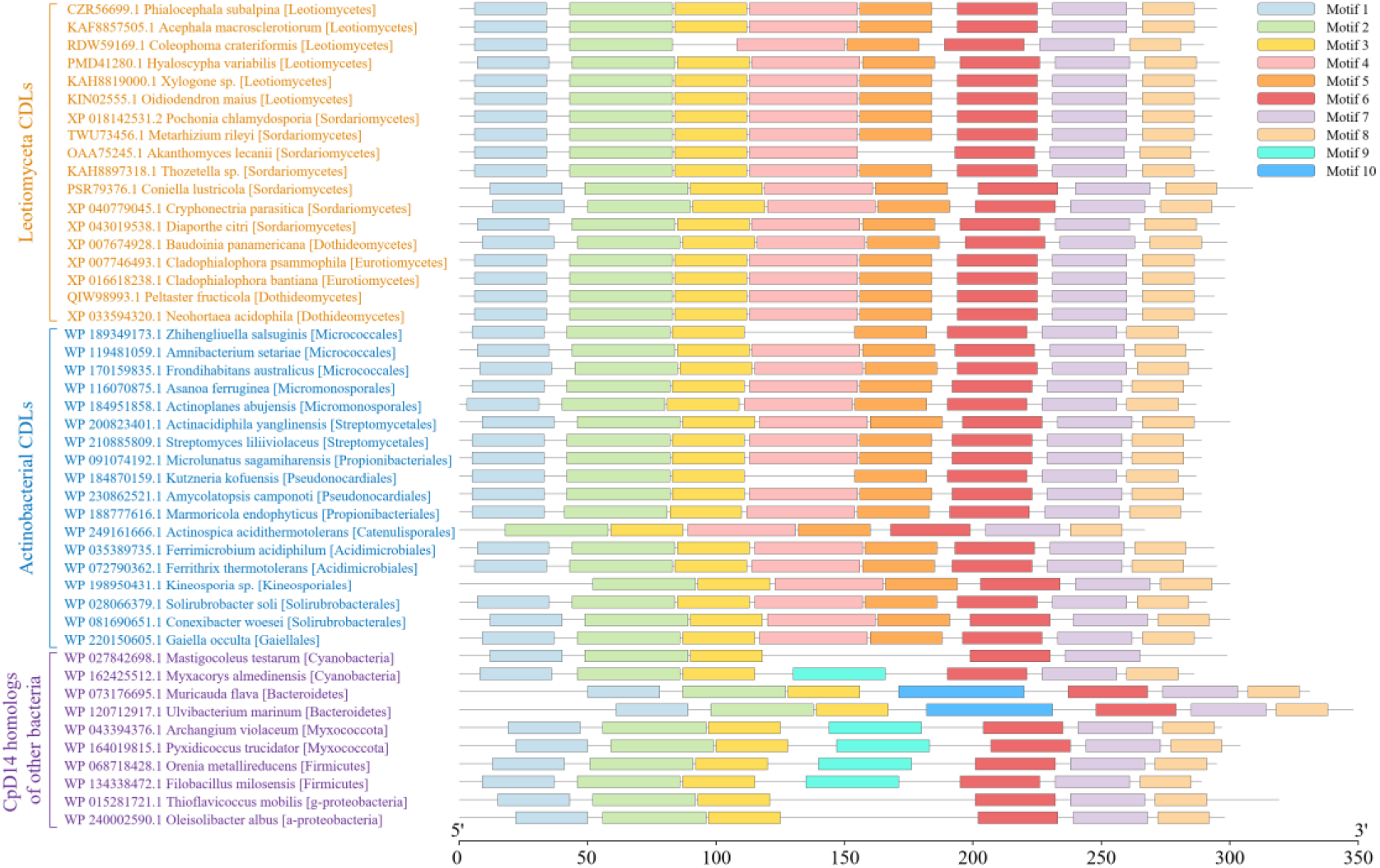
Motif conservation between actinobacterial and Leotiomyceta CDLs. Conserved motifs were identified using MEME suite (E-value threshold was set to 1e-8). CpD14 homologs from other bacteria were included for comparisons.

**Fig. S3.**
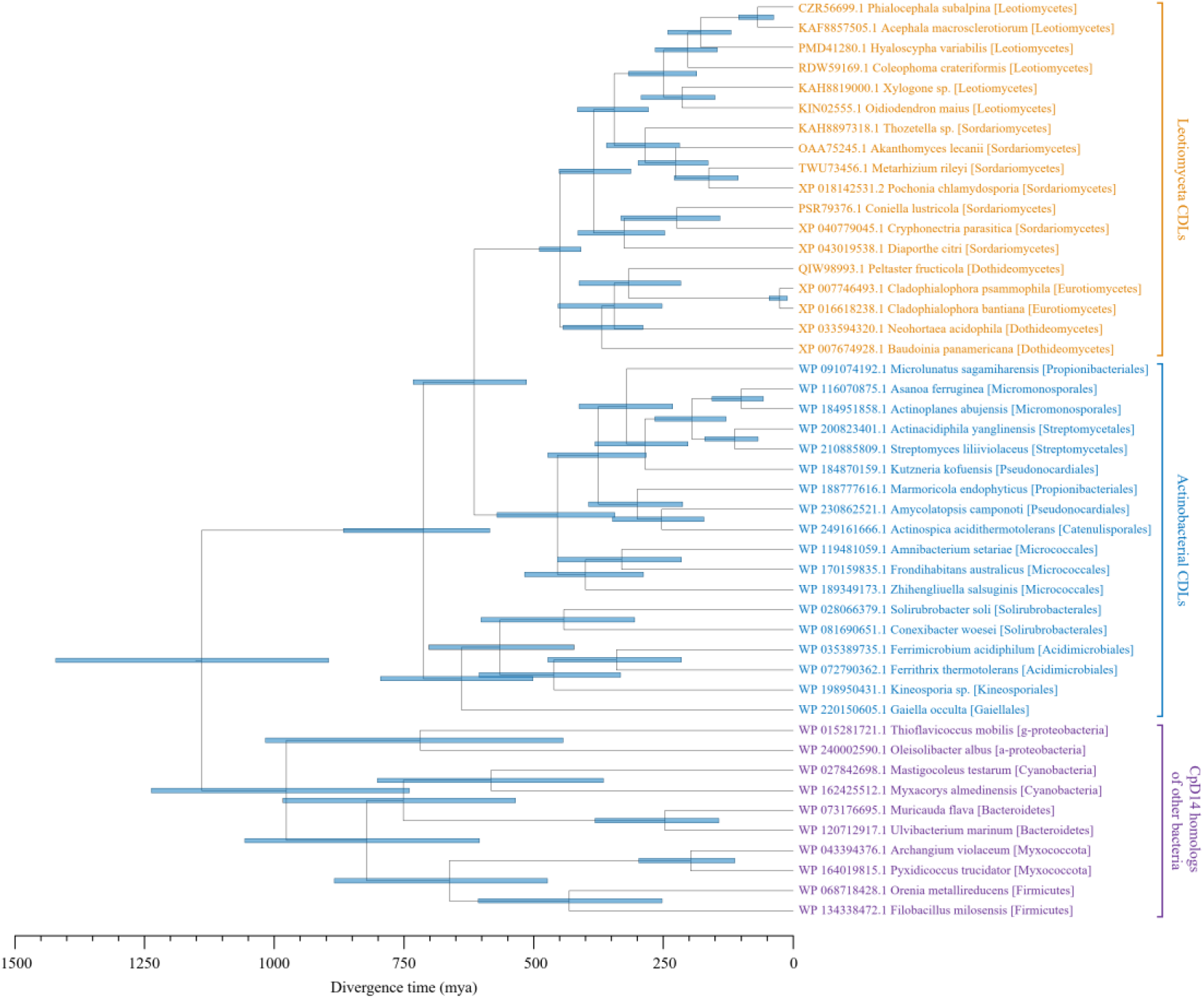
Divergence time estimation of actinobacterial and Leotiomyceta CDLs. The divergence time estimation was performed using BEAST v2.7.4 (62), and node time of Leotiomyceta was set as 450 mya (mean time, with a sigma value of 20 mya) to calibrate the tree according to previous studies (46).

**Fig. S4.**
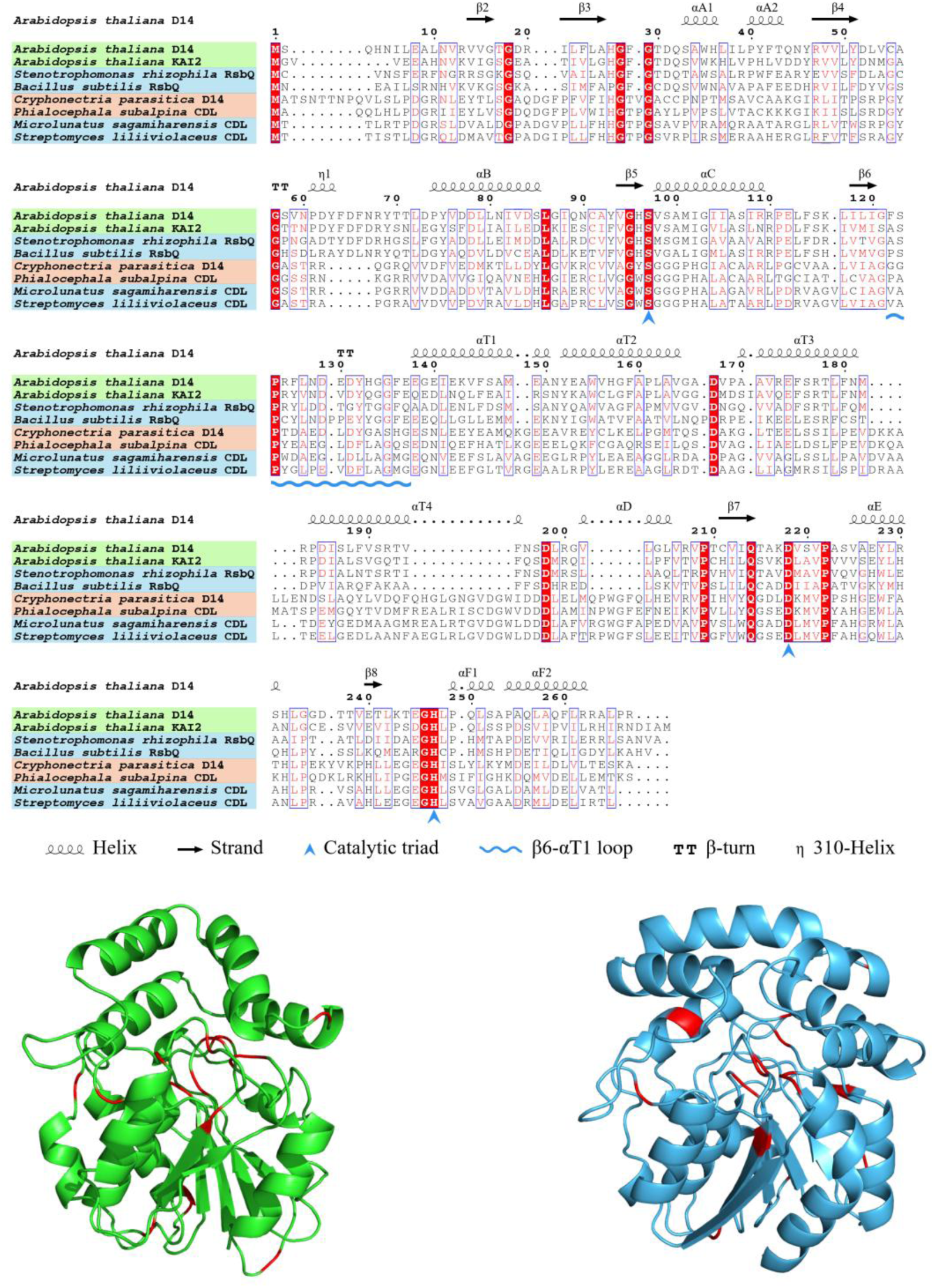
CDLs and DKRs share identical residues for the catalytic triad and several critical folding sites. In the first column, plant, fungal, and bacterial proteins are highlighted with a light green, light brown, and light blue frame, respectively. Secondary structural annotations were based on *Arabidopsis thaliana* D14 (71) (AtD14, PDB ID 4IH4). The tertiary structures of AtD14 (in green) and MsCDL (in blue) are displayed below, and identical residues of the above alignment diagram are marked in red. Critical folding sites refer to 3_10_-helices, α-turns, and β-turns where α-helices or β-strands change direction.

**Fig. S5.**
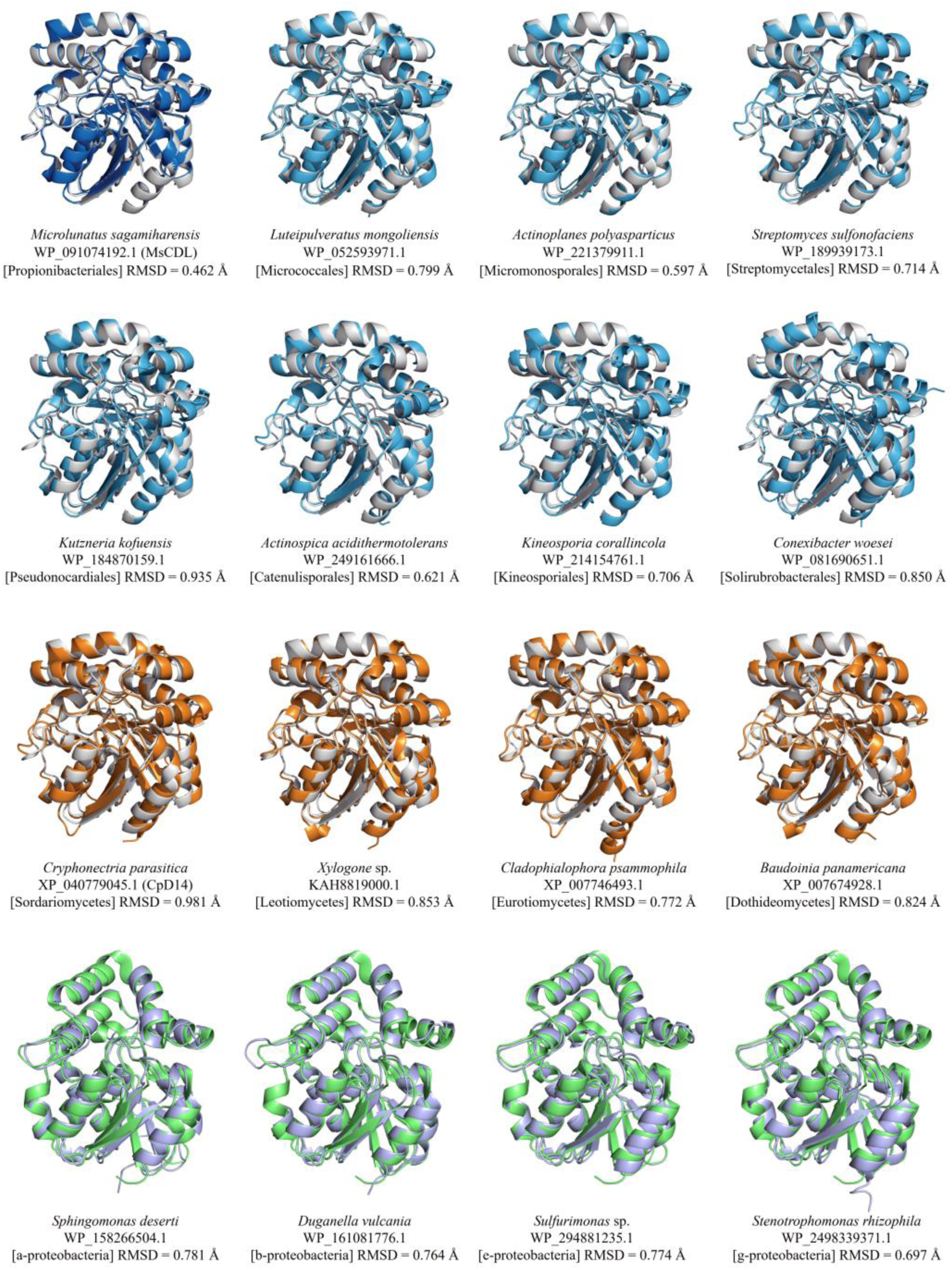
Structural comparisons between the experimentally determined and AlphaFold2 predicted structures of CDLs and DKRs. The AlphaFold2 (30) predicted structures of actinobacterial CDLs (in light blue) and Leotiomyceta CDLs (in brown) were aligned to the experimentally determined MsCDL (in gray), and the predicted proteobacterial RsbQs (in purple) were aligned to OsD14 (in green). The predicted structure of MsCDL (in dark blue) was used as an assessment term to evaluate the accuracy of AlphaFold2 predictions. The RMSD values were calculated with PyMOL.

**Fig. S6.**
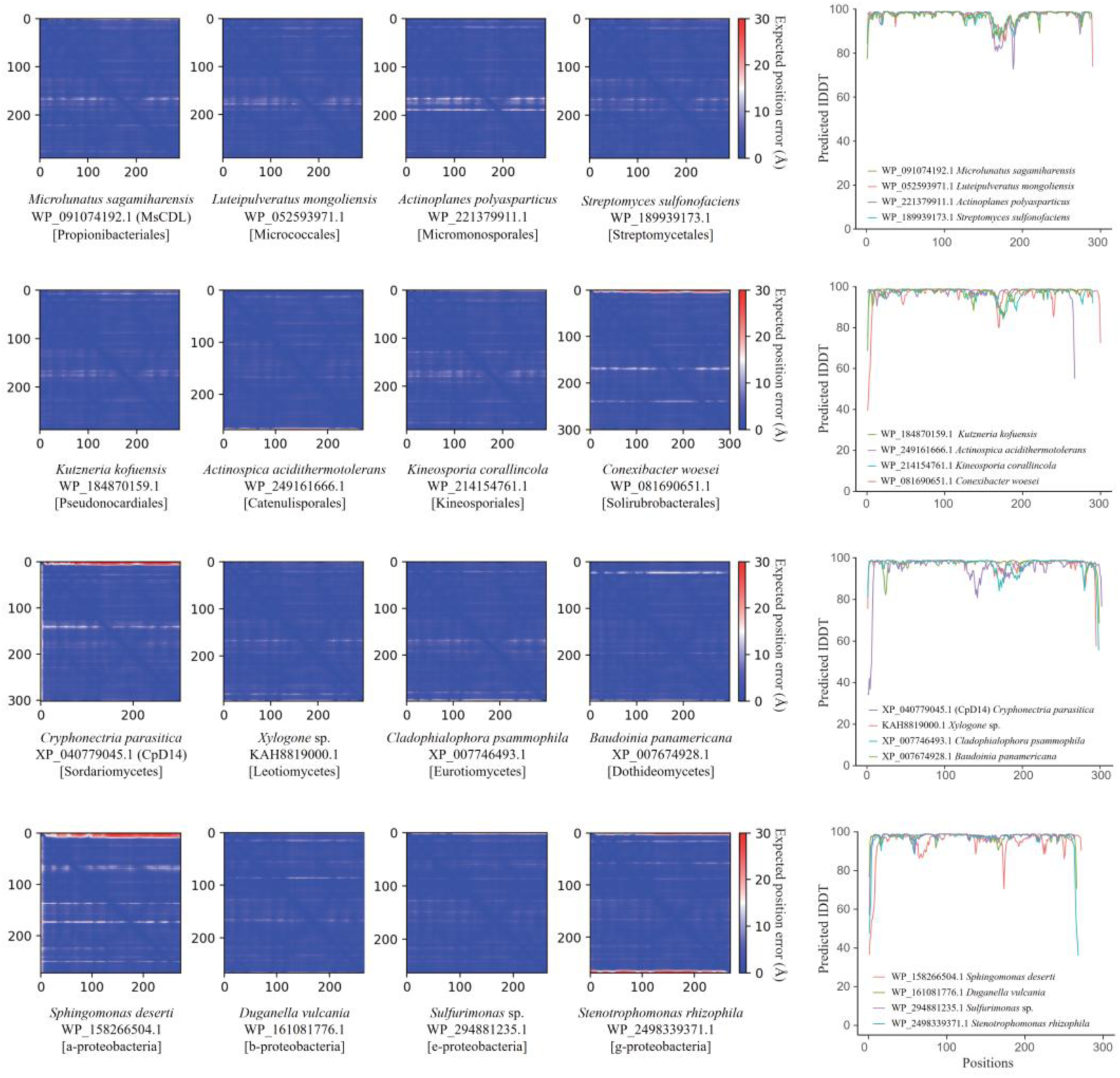
Accuracy and reliability assessment for predicted CDL and RsbQ structures in Fig. S5. The predicted aligned error (PAE) matrixes with a predicted TM (pTM) score are displayed on the left, and the results of predicted local-distance difference test (pLDDT) are displayed on the right. The predicted tertiary structures of CDLs all have a high pLDDT score (average larger than 95) and pTM score (larger than 0.9), suggesting that they are predicted with high accuracy and reliability.

**Fig. S7.**
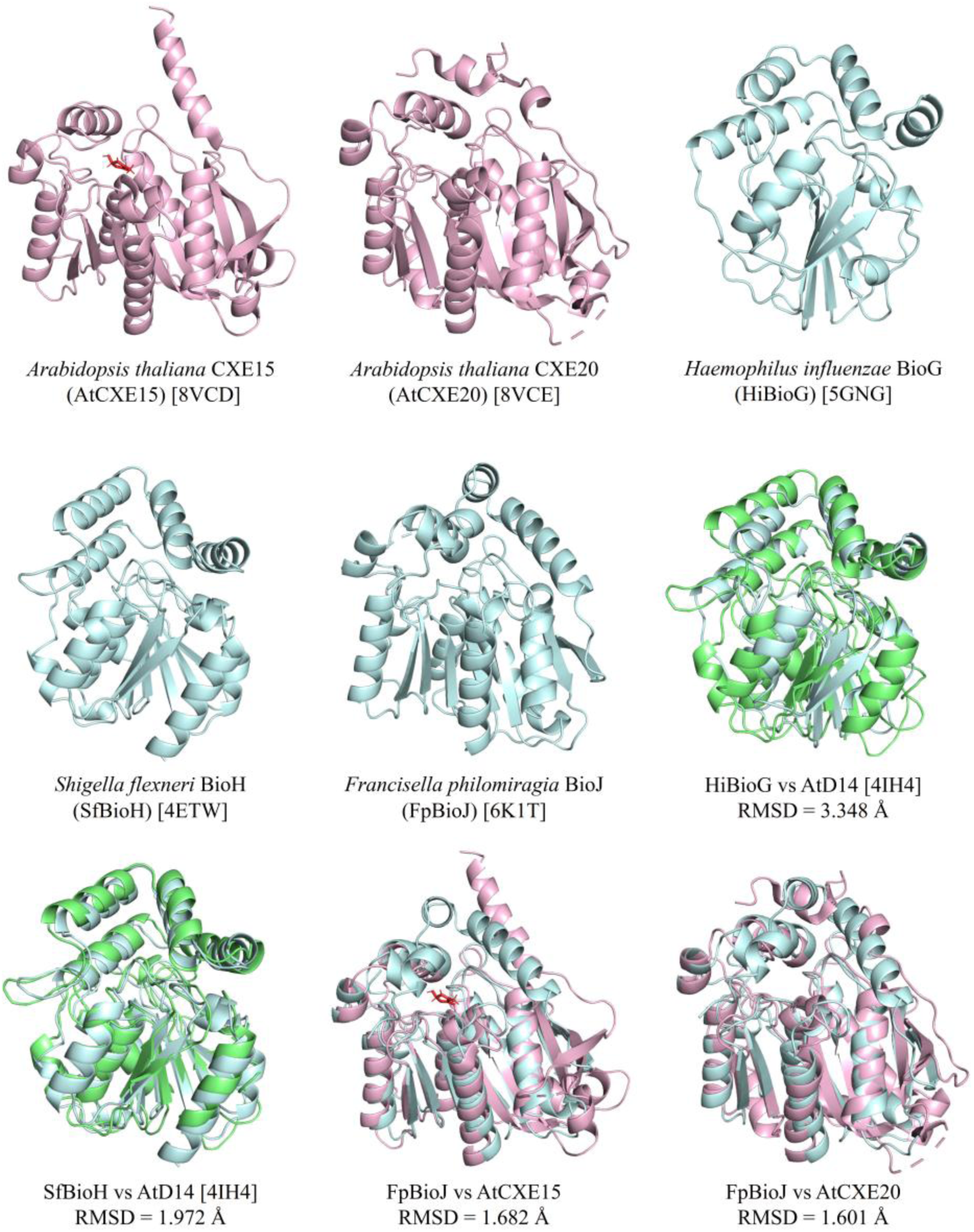
Structural comparisons of plant and bacterial carboxylesterases (CXEs). The structures of *Arabidopsis thaliana* CXEs (AtCXEs, in pink), AtD14 (in green), and bacterial pimeloyl-ACP methyl ester CXEs (in light blue) were downloaded from the PDB (PDB IDs noted in the square brackets).

**Fig. S8.**
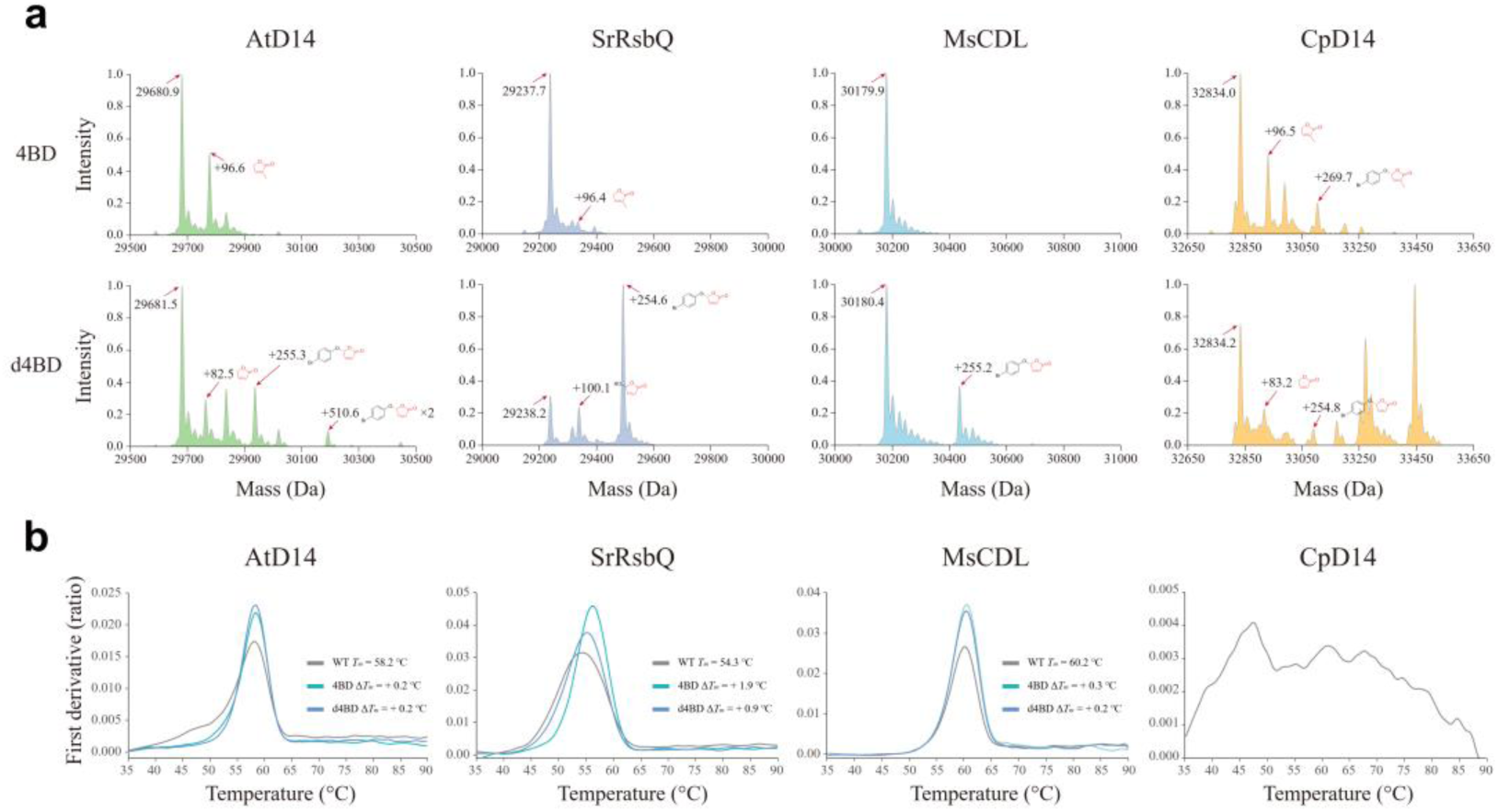
Binding activity of DKRs and CDLs toward 4BD and d4BD. a, Deconvoluted intact mass spectra of DKRs and CDLs after 1 h incubation with 4BD and d4BD, respectively. The peaks of native proteins and their putative bound states are labeled with molecular weight values. **b,** Thermal shift assays. The melting temperature curves represent the means of 3 technical replicates. The melting temperature (*T*_m_) of protein and its variations (Δ*T*_m_) after 30 min incubation with substrates are shown on the right of each group.

**Fig. S9.**
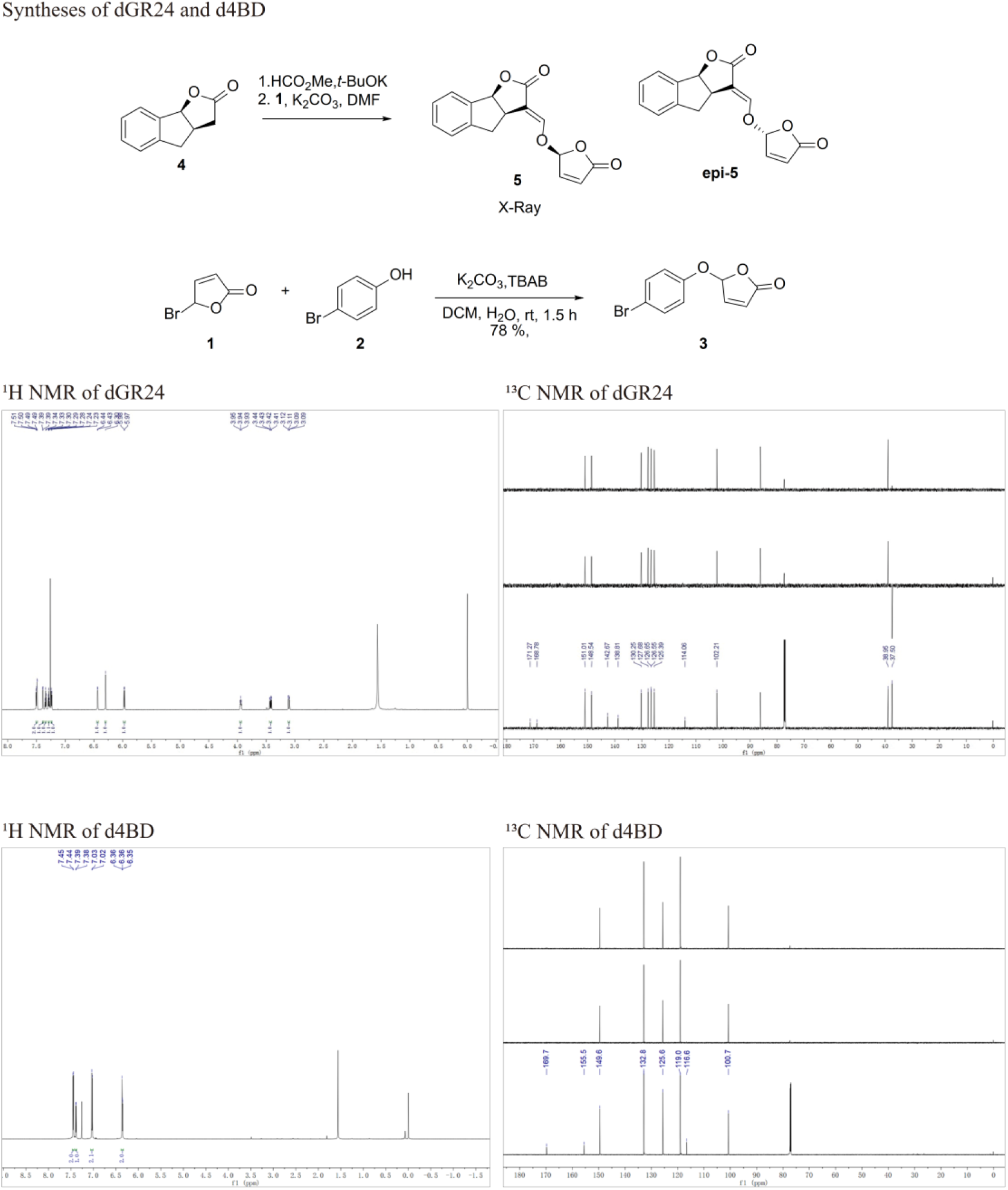
The syntheses and NMR analyses of dGR24 and d4BD. To a solution of (±)-**4** (64 mg, 0.37 mmol, 1.0 equiv) in HCO_2_Me (1 mL) was added *t*-BuOK (207 mg 1.85 mmol, 5.0 equiv) under atmosphere of nitrogen at 0 °C. The reaction mixture was stirred overnight at room temperature. The reaction was quenched with 1.0 N aqueous HCl solution (2.5 mL), and the aqueous phase was extracted with EtOAc (3 × 5 mL). The combined organic phase was washed with brine, dried over Na_2_SO_4_, filtered and concentrated *in vacuo*. To a solution of the above residue in DMF (2 mL) was added K_2_CO_3_ (77 mg, 0.56 mmol, 1.5 equiv) and bromobutenolide **1** (90.5 mg, 0.56 mmol, 1.5 equiv) in DMF (1 mL) at 0 °C. The solution was stirred overnight at room temperature. The mixture was then quenched with saturated *aq.* NH_4_Cl (5 mL) and extracted with EtOAc (3 × 5 mL). The combined organic phase was washed with brine, dried over Na_2_SO_4_, filtered and concentrated *in vacuo*. The crude residue was purified by column chromatography (petroleum ether/EtOAc 2:1) to give the product **5** (35.7 mg, 34% yield) and ***epi*-5** (38.9 mg, 37% yield). NMR data of dGR24: ^1^H NMR (800 MHz, CDCl_3_) *δ* 3.10 (dd, *J* = 16.8, 2.7 Hz, 1H), 3.42 (dd, *J* = 16.8, 9.3 Hz, 1H), 3.94 (m, 1H), 5.97 (d, *J* = 7.8 Hz, 1H), 6.30 (s, 1H), 6.44 (d, *J* = 5.6 Hz, 1H), 7.24 (d, *J* = 7.6 Hz, 1H), 7.29 (t, *J* = 7.4 Hz, 1H), 7.34 (t, *J* = 7.4 Hz, 1H), 7.39 (d, *J* = 5.6 Hz, 1H), 7.29 (d, *J* = 2.4 Hz, 1H), 7.50 (d, *J* = 7.6 Hz, 1H). ^13^C NMR (200 MHz, CDCl_3_) *δ* 171.3, 168.3, 151.0, 148.5, 142.7, 138.8, 130.2, 127.7, 126.6, 126.6, 125.4, 114.1, 102.2, 86.1, 39.0, 37.5. HRESIMS *m/z* 307.0582 [M + Na]^+^ (calcd for C_16_H_12_O_5_Na, 307.0582). To a solution of alcohol **2** (34.6 mg, 0.2 mmol, 1.0 equiv) in DCM (2 mL) were added tetrabutyl ammonium bromide (TBAB, 6.4 mg, 0.02 mmol, 0.1 equiv) and K_2_CO_3_ (41.4 mg, 0.3 mmol, 1.5 equiv) in H_2_O (2 mL) sequentially at room temperature. After vigorous stirring for 10 min at room temperature, a solution of bromobutenolide **1** (39.1 mg, 0.24 mmol, 1.2 equiv) in DCM (2 mL) was added dropwise. The two phase mixture was stirred vigorously for another 1.5 h, the organic phase was separated, and the aqueous phase was washed with DCM (3 × 5 mL). The combined organic phase was washed with brine, dried over Na_2_SO_4_, filtered and concentrated *in vacuo*. The crude residue was purified by column chromatography (petroleum ether/EtOAc 15:1) to give the product **3** (39.8 mg, 78% yield). NMR data of d4BD: ^1^H NMR (600 MHz, CDCl_3_) *δ* 6.35 (d, *J* = 5.7 Hz, 1H), 6.36 (s, 1H), 7.03 (dd, *J* = 8.7, 2.5 Hz, 2H), 7.39 (d, *J* = 5.7 Hz, 1H), 7.45 (dd, *J* = 8.7, 2.5 Hz, 2H). ^13^C NMR (150 MHz, CDCl_3_) *δ* 169.7, 155.5, 149.6, 132.8, 125.6, 119.0, 116.6, 100.7. HREIMS *m/z* 253.9574 [M]^+^ (calcd for C_10_H_7_O_3_Br, 253.9573).

**Table S1.** X-ray data collection and refinement statistics of MsCDL.

| Crystal | 1R670 |
| --- | --- |
| Data collection |  |
| Wavelength (Å) | 0.95373 |
| Space group | P41212 |
| Unit cell dimensions |  |
| *a*, *b*, *c* (Å) | 125.21, 125.21, 79.73 |
| *α*, *β*, *γ* (°) | 90.00, 90.00, 90.00 |
| Resolution (Å) | 49.24-1.90 (1.94-1.90) |
| No. of observations | 1270417 (84647) |
| No. Unique | 50439 (3188) |
| Completeness (%) | 100.0 (100.0) |
| CC1/2 | 0.999 (0.918) |
| Redundancy | 25.2 (26.6) |
| *R*<sub>merge</sub> (*R*<sub>sym</sub>)(%) | 13.1 (98.6) |
| *R*<sub>meas</sub> (%) | 13.4 (100.5) |
| *I*/*σ* (I) | 21.6 (4.4) |
| Refinement |  |
| Resolution (Å) | 49.29-1.90 (1.95-1.90) |
| No. of reflections | 47850 (3472) |
| *R*<sub>work</sub>/*R*<sub>free</sub> (%) | 14.40/17.94 |
| No. of atoms |  |
| Protein | 4281 |
| Ion | 11 |
| Water | 634 |
| Others | 77 |
| *B*-factors |  |
| Protein | 24.080 |
| Ion | 69.828 |
| Water | 36.299 |
| Others | 48.404 |
| R.m.s deviations |  |
| Bond lengths (Å) | 0.012 |
| Bond angles (°) | 1.636 |
| Ramachandran plot (%) |  |
| Most favored | 97.02 |
| Allowed | 2.45 |
| Disallowed | 0.53 |

\* Statistics for the highest-resolution shell are shown in parentheses

